# *Mycobacterium leprae* and host immune transcriptomic signatures for reactional states in leprosy

**DOI:** 10.1101/2022.07.25.501496

**Authors:** Madhusmita Das, Diana David, Ilse Horo, Anouk Van Hooij, Maria Tió Coma, Annemieke Geluk, Sundeep Chaitanya Vedithi

## Abstract

**Background:** *Mycobacterium leprae* transcriptomic and human host immune gene expression signatures that demonstrate a plausible association with type I (T1R) and type II reactions (T2R) aid in early diagnosis, prevention of nerve damage and consequent demyelinating neuropathy in leprosy. The aim of the study is to identify *M. leprae* and host-associated gene-expression signatures that are associated with reactional states in leprosy.

**Methods:** The differentially expressed genes from the whole transcriptome of *M. leprae* were determined using genome-wide hybridization arrays with RNA extracted from skin biopsies of 20 T1R, 20 T2R and 20 non reactional controls (NR). Additionally, human immune gene-expressions were profiled using RT2-PCR profiler arrays and real-time qPCRs.

**Results:** The RNA quality was optimal in 16 NR, 18 T1R and 19 T2R samples. Whole transcriptome expression array of these samples revealed significant overexpression of the genes that encode integral and intrinsic membrane proteins, hydrolases and oxidoreductases. In T1R lesional skin biopsy specimens, the top 10 over significantly upregulated genes are ML2064, ML1271, ML1960, ML122, ML2498, ML1996, ML2388, ML0429, ML2030 and ML0224 in comparison to NR. In T2R, genes ML2498, ML1526, ML0394, ML1960, ML2388, ML0429, ML0281, ML1847, ML1618 and ML1271 were significantly upregulated. We noted ML2664 was significantly upregulated in T1R and repressed in T2R. Conversely, we haven’t noted any genes upregulated in T2R and repressed in T1R. In both T1R and T2R, ML2388 was significantly overexpressed. This gene encodes a probable membrane protein and epitope prediction using Bepipred −2.0 revealed a distinct B-cell epitope. Overexpression of ML2388 was noted consistently across the reaction samples.

From the host immune gene expression profiles, genes for CXCL9, CXCL2, CD40LG, IL17A and CXCL11 were upregulated in T1R when compared to the NR. In T2R, CXCL10, CXCL11, CXCL9, CXCL2 and CD40LG were upregulated.

**Conclusion:** A gene set signature involving bacterial genes ML2388, ML2664, and host immune genes CXCL10 and IL-17A can be transcriptomic markers for reactional states in leprosy.

## INTRODUCTION

*Mycobacterium leprae* (*M. leprae*), the causative bacillus for leprosy, continues to infect endemic populations in tropical countries, with approximately 200,000 new cases of leprosy emerging each year globally. *M. leprae* infects the skin and the peripheral nerves causing skin lesions with loss of sensation resulting from demyelinating neuropathy as the bacilli infect the Schwann cells of the axonal myelin in the peripheral neurons (1). Nerve damage in leprosy is mediated by *M. leprae* infection of the Schwann cells as well as exacerbated immune responses in the human host. Leprosy is manifested with a complex host immunological profile that classifies the disease into a cell-mediated immunity (CMI) - dominated tuberculoid pole (TT) and the humoral immune (HI) response - regulated lepromatous pole (LL)(2). Both these poles are separated by three borderline intermediary groups that gradient from CMI towards the HI. These include the borderline-tuberculoid (BT), mid-borderline (BB) and marginal lepromatous forms (BL).(3)

About 30-40% of leprosy infected individuals in the borderline forms and rarely in the polar states manifest delayed-type hypersensitivity reactions, the type 1 reaction also known as reversal reaction and the type 2 reaction known as Erythema Nodusum Leprosum (ENL) (4,5). These inflammatory responses can occur before, during and after the treatment with multidrug therapy (MDT) and are managed by immunomodulatory drugs in high doses that often contribute to morbidity. Reactional states are a significant cause of nerve damage and associated disability in leprosy. Early detection of reactional episodes can facilitate prophylactic treatment interventions that minimize the risk of nerve damage(6–8).

Predictive genomic, transcriptomic and host immune biomarkers can play a critical role in detecting subclinical nerve damage and determining factors that trigger reactional states in leprosy. In leprosy endemic tropical countries, it is often challenging to characterize an individual’s immune background due to varied antigenic exposure(9). Thus, attributing specific immune responses (cytokine and antibody quantities or human immune gene expression signatures) alone to the onset of reactional states or leprosy per se may offer limited applicability in developing effective diagnostics for these *M. leprae* specific immune exacerbations in leprosy(10).

A correlative gene expression signature that originates from both *M. leprae* and human host immune system provides comprehensive predictive and prognostic information for determining the onset of these inflammatory responses. In this study, we conducted a cross sectional analysis to quantify relative abundance of *M. leprae* and host immune gene transcripts in localized skin lesions of leprosy cases with type I and type II reactions. A gene expression signature that demonstrates significant association with reactional states has been determined. Follow up studies are warranted to validate these expression signatures in a longitudinal cohort(11).

## MATERIALS AND METHOD

### Sample Size

A total of 60 newly diagnosed untreated leprosy cases were recruited at the outpatient department of Schieffelin Institute of Health Research and Leprosy Centre in Karigiri, India. Following institutional ethical clearance, informed and written consent for participation was obtained from each subject prior to recruitment in the study following the ethical guidelines as laid down by the Indian Council of Medical Research. The sample was stratified as 20 with Type 1 Reaction, 20 with Type 2 reaction and 20 without any reaction. Post clinical examination, 5mm x 5mm excisional skin biopsies from skin lesions were collected of subjects for all the study groups. Clinical details of the sample were provided in table 1.

**Table1:** Clinical and Demographic characteristics of the study sample.

| SL NO. | Sample ID | Reactional status | Age | Gender | WHO Classification | RJ Classification | Bacteriological Index |
| --- | --- | --- | --- | --- | --- | --- | --- |
| 1 | LRI 001 | T2R | 47 | Male | MB | LL | 4.75 |
| 2 | LRI 002 | NR | 40 | Male | MB | HISTOID | 2.75 |
| 3 | LRI 003 | NR | 33 | Female | MB | BT | 0.5 |
| 4 | LRI 004 | NR | 50 | Male | MB | BT | 0 |
| 5 | LRI 005 | NR | 52 | Male | MB | LL | 3.25 |
| 6 | LRI 006 | T2R | 48 | Male | MB | LL | 5 |
| 7 | LRI 007 | NR | 15 | Male | MB | BL | 3.5 |
| 8 | LRI 008 | NR | 29 | Male | MB | LL | 4 |
| 9 | LRI 009 | NR | 32 | Female | MB | LL | 4.25 |
| 10 | LRI 010 | NR | 52 | Male | MB | BL | 3 |
| 11 | LRI 011 | NR | 40 | Male | PB | TT | 0 |
| 12 | LRI 012 | T1R | 47 | Male | MB | BL | 2 |
| 13 | LRI 013 | NR | 50 | Male | MB | BT | 0 |
| 14 | LRI 014 | NR | 35 | Female | MB | LL | 4 |
| 15 | LRI 015 | NR | 17 | Female | MB | LL | 3 |
| 16 | LRI 016 | NR | 60 | Male | MB | LL | 4 |
| 17 | LRI 017 | NR | 42 | Male | MB | BT | 0 |
| 18 | LRI 018 | T1R | 35 | Female | MB | LL | 4.5 |
| 19 | LRI 019 | NR | 35 | Male | MB | BL | 3.25 |
| 20 | LRI 020 | NR | 52 | Male | MB | LL | 4 |
| 21 | LRI 021 | NR | 26 | Male | MB | LL | 0 |
| 22 | LRI 022 | NR | 25 | Male | MB | BT | 0 |
| 23 | LRI 023 | T1R | 29 | Male | MB | BT | 0 |
| 24 | LRI 024 | T1R | 33 | Male | MB | BT | 0 |
| 25 | LRI 025 | T1R | 30 | Female | MB | BT | 0 |
| 26 | LRI 026 | NR | 27 | Female | PB | BT | 0 |
| 27 | LRI 027 | NR | 35 | Male | MB | LL | 2.5 |
| 28 | LRI 028 | T1R | 35 | Male | MB | LL | 4.75 |
| 29 | LRI 029 | T2R | 58 | Male | MB | LL | 3.25 |
| 30 | LRI 030 | T1R | 18 | Female | MB | LL | 1.5 |
| 31 | LRI 031 | T1R | 42 | Male | MB | LL | 3 |
| 32 | LRI 032 | T2R | 35 | Female | MB | BL | 4.25 |
| 33 | LRI 033 | T1R | 20 | Male | MB | BL | 2.25 |
| 34 | LRI 034 | T1R |  | Male | MB | BB | 0 |
| 35 | LRI 035 | T1R | 29 | Female | MB | BT | 1 |
| 36 | LRI 036 | T1R | 30 | Female | MB | BL | 4 |
| 37 | LRI 037 | T1R | 43 | Male | MB | BL | 0.25 |
| 38 | LRI 038 | T2R | 27 | Male | MB | BL | 4 |
| 39 | LRI 039 | T1R | 61 | Male | MB | BL | 1 |
| 40 | LRI 040 | T1R | 42 | Male | MB | BT | 0 |
| 41 | LRI 041 | T1R | 32 | Male | PB | BT | 0 |
| 42 | LRI 042 | T2R | 35 | Male | MB | LL | 4 |
| 43 | LRI 043 | T1R | 36 | Female | MB | BT | 0 |
| 44 | LRI 044 | T1R | 55 | Male | MB | BL | 0 |
| 45 | LRI 045 | T1R | 40 | Female | MB | LL | 2.75 |
| 46 | LRI 046 | T2R | 82 | Male | MB | LL | 3 |
| 47 | LRI 047 | T1R | 52 | Male | MB | BL | 2.25 |
| 48 | LRI 048 | T2R | 24 | Female | MB | LL | 0 |
| 49 | LRI 049 | T2R | 4 | Female | MB | LL | 3 |
| 50 | LRI 050 | T2R | 21 | Female | MB | LL | 3 |
| 51 | LRI 051 | T2R | 32 | Male | MB | LL | 3.5 |
| 52 | LRI 052 | T2R | 35 | Male | MB | LL | 4 |
| 53 | LRI 053 | T2R | 24 | Female | MB | LL | 3 |
| 54 | LRI 054 | T2R | 22 | Female | MB | LL | 3 |
| 55 | LRI 055 | T2R | 28 | Male | MB | BL | 2.35 |
| 56 | LRI 056 | T2R | 30 | Female | MB | LL | 3+ |
| 57 | LRI 057 | T2R | 37 | Male | MB | LL | 3.75 |
| 58 | LRI 058 | T2R | 19 | Male | MB | LL | 3 |
| 59 | LRI 059 | T2R | 32 | Male | MB | LL | 3.5 |
| 60 | LRI 060 | T2R | 26 | Male | MB | LL | 6 |

### *M. leprae* Whole Transcriptome Hybridization Arrays

Total RNA was extracted following the Trizol protocol (Qiagen RNeasy Lipi Tissue kit - Cat#74804) and bacterial RNA was enriched in the samples. The quality of RNA was estimated using BioAnalyzer 2100 (Agilent Technologies) followed by labelling, reverse transcription, amplification and hybridization to the arrays. Based on the BioAnalyzer reports, 16 NR, 18 T1R and 19 T2R samples were found suitable for whole transcriptome hybridizations.

A 2×400K gene expression array (whole-genome tiling array) was designed with the probes having 60-mer oligonucleotides tiling every 10bp of the genome sequence of *M. leprae* (NC_011896.1). The array comprised 420288 features which include probes and Agilent controls. The samples for gene expression were labelled using the Agilent Quick-Amp labelling Kit (p/n5190-0442). 500ng each of total RNA was reverse transcribed at 40°C using oligo dT primer tagged to a T7 polymerase promoter and converted to double-stranded cDNA. Synthesized double-stranded cDNA were used as templates for cRNA generation. cRNA was generated by *in-vitro* transcription and the dye Cy3 CTP (Agilent) was incorporated during this step. The cDNA synthesis and *in-vitro* transcription steps were carried out at 40°C. Labelled cRNA was cleaned up using Qiagen RNeasy columns (Qiagen, Cat No: 74106) and quality was assessed for yields and specific activity using the Nanodrop ND-1000. The hybridized slides were scanned on a G2600D scanner (Agilent Technologies). The data thus acquired is analysed using GeneSpring GX Version 12.1 software. Data were normalized and a fold difference in expression was noted from 359,922 probes which include sense and antisense orientations of 179,961 probes. The differentially expressing *M. leprae* genomic regions between type 1, type 2 reactions and non-reactional cases were noted. All the samples were performed in technical replicates to validate the observations and microarray data corresponding to 359,922 probes for each of the samples (Fig 1). The data as well as the array design was uploaded to the Gene Expression Omnibus (GEO) Repository of the National Center for Biotechnology Information (NCBI) with the accession numbers: GSE85948 and GPL22363.

**Fig-1:**
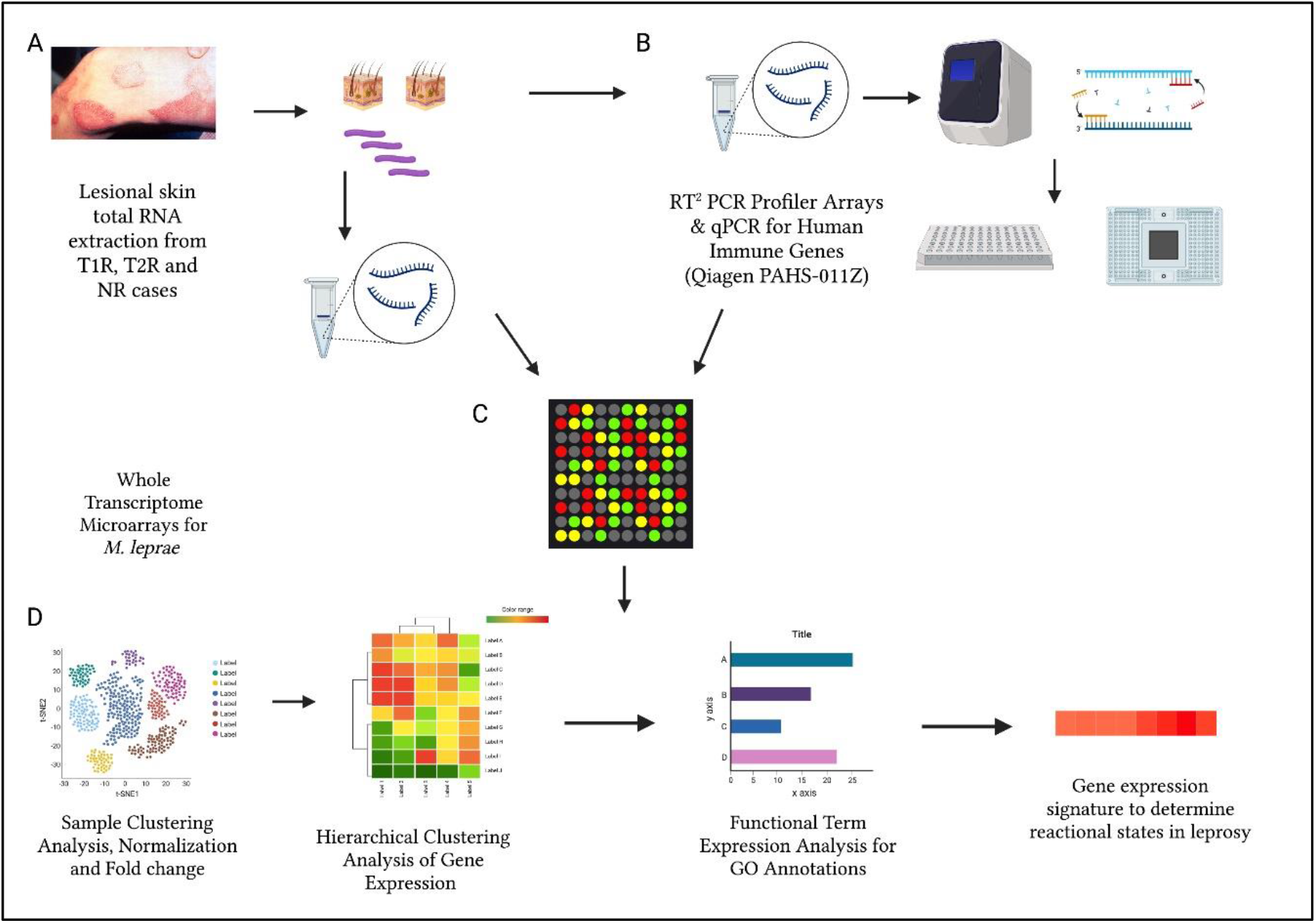
The schematic representation for the cross-sectional analysis of *M. leprae* transcriptome (A), human immune gene expression (B) and corresponding circulatory levels of cytokines, interleukins and chemokines (C) to derive a gene expression signature for reactional states of leprosy (D). (Created with BioRender.com.)

The 75th percentile ranking was used to normalize the probe intensities. The fold difference in expression was noted by subtracting the gene intensities of reactional samples from that of non-reactional samples in each experiment using the geometric mean of the technical replicates. These fold changes were log-transformed to base 2 and volcano plots were generated to identify differentially expressed genes (DEGs). The fold change of >=0.6 was considered as upregulated and <=-0.6 was considered as down-regulated. The Benjamin Hochberg adjusted P values were represented as −log_10_ (P value)(12).

### RT^2^ PCR Profiler Arrays

RT^2^ PCR profiler arrays (Qiagen Inc, USA) for human inflammatory cytokines and receptors (PAHS-011Z) were used to quantitate expression levels of 96 human immune genes in the lesional skin RNA samples across the study group. Each catalogued RT^2^Profiler PCR Array contains a list of the human inflammatory cytokines and receptors genes as well as five housekeeping (reference) genes on the array. In addition, each array contains a panel of proprietary controls to monitor genomic DNA contamination (GDC) as well as the first strand synthesis (RTC) and real-time PCR efficiency (PPC). The list of genes was provided in Qiagen array Cat. no. PAHS-011Z. Total RNA was isolated from skin biopsy specimens using RNeasy kit (Qiagen Cat No: Cat. No. / ID: 74104) according to the manufacturer’s instructions. RNA quality was determined using a Nanodrop and was reverse transcribed using a QuantiTect Reverse Transcription Kit (Cat No: Cat. No. / ID: 205311). The cDNA was used on the real-time RT^2^ Profiler PCR Array (Cat. no. PAHS-011Z) in combination with RT^2^SYBR Green qPCR Mastermix (Qiagen Cat. no. 330529). Fold-change values greater than one indicates a positive- or an up-regulation, and the fold-regulation is equal to the fold-change. Fold-change values less than one indicate a negative or down-regulation, and the fold-regulation is the negative inverse of the fold-change. The *p-values* were calculated based on a student’s t-test of the replicate 2^(-Delta Ct)^ values for each gene in the control group and treatment groups.

### Multiplex qPCR

RNA extracted from lesional skin biopsy specimens (NR=16; T1R=16; T2R=9) was used in the qPCR assays. The genes GNLY, CD8A, CXCL10, IFI6, IL10, PRF1, CCL2, FCGR1B, OAS1, IFI44, and CTLA4 have been amplified using the conditions described elsewhere (Table 4)(13). The data of the qPCR were analyzed using Thermo Fisher Cloud and GraphPad Prism version 8.0.1. Samples were analyzed in duplicates and ΔCts were calculated using GAPDH as the reference gene. Mann-Whitney U test was performed to determine if differences in gene expression is statistically significant.

**Table 2:** The enriched GO terms in T1R and T2R groups across the sample.

| GO Term | GO Type | Reaction Type | Count | p-value |
| --- | --- | --- | --- | --- |
| GO:0016021~integral component of membrane | Cellular Localization | T1R | 44 | 0.000041 |
| GO:0031224~intrinsic component of membrane | Cellular Localization | T1R | 44 | 0.000045 |
| GO:0044425~membrane part | Cellular Localization | T1R | 44 | 0.000060 |
| GO:0016020~membrane | Cellular Localization | T1R | 44 | 0.000197 |
| GO:0006351~transcription, DNA-templated | Biological Process | T1R | 12 | 0.021228 |
| GO:0097659~nucleic acid-templated transcription | Biological Process | T1R | 13 | 0.011823 |
| GO:1903506~regulation of nucleic acid-templated transcription | Biological Process | T1R | 9 | 0.005321 |
| GO:0006355~regulation of transcription, DNA-templated | Biological Process | T1R | 9 | 0.005321 |
| GO:2001141~regulation of RNA biosynthetic process | Biological Process | T1R | 9 | 0.005321 |
| GO:0051252~regulation of RNA metabolic process | Biological Process | T1R | 9 | 0.006193 |
| GO:0032774~RNA biosynthetic process | Biological Process | T1R | 13 | 0.017436 |
| GO:0019219~regulation of nucleobase-containing compound metabolic process | Biological Process | T1R | 9 | 0.008264 |
| GO:0010468~regulation of gene expression | Biological Process | T1R | 9 | 0.009478 |
| GO:2000112~regulation of cellular macromolecule biosynthetic process | Biological Process | T1R | 9 | 0.010821 |
| GO:0031326~regulation of cellular biosynthetic process | Biological Process | T1R | 9 | 0.010821 |
| GO:0009889~regulation of biosynthetic process | Biological Process | T1R | 9 | 0.010821 |
| GO:0010556~regulation of macromolecule biosynthetic process | Biological Process | T1R | 9 | 0.010821 |
| GO:0051171~regulation of nitrogen compound metabolic process | Biological Process | T1R | 9 | 0.013930 |
| GO:0080090~regulation of primary metabolic process | Biological Process | T1R | 9 | 0.015711 |
| GO:0060255~regulation of macromolecule metabolic process | Biological Process | T1R | 9 | 0.015711 |
| GO:0006629~lipid metabolic process | Biological Process | T1R | 12 | 0.029266 |
| GO:0031323~regulation of cellular metabolic process | Biological Process | T1R | 9 | 0.017654 |
| GO:0019222~regulation of metabolic process | Biological Process | T1R | 9 | 0.017654 |
| GO:0003677~DNA binding | Molecular Function | T1R | 14 | 0.048031 |
| GO:0006810~transport | Biological Process | T1R | 15 | 0.031141 |
| GO:0051234~establishment of localization | Biological Process | T1R | 15 | 0.031141 |
| GO:0051179~localization | Biological Process | T1R | 15 | 0.031141 |
| GO:0042221~response to chemical | Biological Process | T1R | 6 | 0.012239 |
| GO:0071840~cellular component organization or biogenesis | Biological Process | T1R | 13 | 0.037976 |
| GO:0046677~response to antibiotic | Biological Process | T1R | 5 | 0.021659 |
| GO:0022607~cellular component assembly | Biological Process | T1R | 5 | 0.044536 |
| GO:0050896~response to stimulus | Biological Process | T1R | 11 | 0.035335 |
| GO:0016787~hydrolase activity | Molecular Function | T1R | 39 | 0.045506 |
| GO:0016021~integral component of membrane | Cellular Localization | T2R | 45 | 0.008044 |
| GO:0031224~intrinsic component of membrane | Cellular Localization | T2R | 45 | 0.008261 |
| GO:0044425~membrane part | Cellular Localization | T2R | 50 | 0.003330 |
| GO:0016020~membrane | Cellular Localization | T2R | 51 | 0.007984 |
| GO:0006629~lipid metabolic process | Biological Process | T2R | 12 | 0.036602 |
| GO:0046488~phosphatidylinositol metabolic process | Biological Process | T2R | 3 | 0.037410 |
| GO:0046486~glycerolipid metabolic process | Biological Process | T2R | 4 | 0.044399 |
| GO:0003677~DNA binding | Molecular Function | T2R | 14 | 0.027195 |
| GO:0006351~transcription, DNA-templated | Biological Process | T2R | 4 | 0.044833 |
| GO:0044085~cellular component biogenesis | Biological Process | T2R | 11 | 0.002047 |
| GO:0071840~cellular component organization or biogenesis | Biological Process | T2R | 13 | 0.002468 |
| GO:0006364~rRNA processing | Biological Process | T2R | 5 | 0.006889 |
| GO:0016072~rRNA metabolic process | Biological Process | T2R | 5 | 0.006889 |
| GO:0016070~RNA metabolic process | Biological Process | T2R | 18 | 0.007569 |
| GO:0006396~RNA processing | Biological Process | T2R | 8 | 0.008240 |
| GO:0022613~ribonucleoprotein complex biogenesis | Biological Process | T2R | 5 | 0.032265 |
| GO:0042254~ribosome biogenesis | Biological Process | T2R | 5 | 0.032265 |
| GO:0023052~signaling | Biological Process | T2R | 4 | 0.034417 |
| GO:0007165~signal transduction | Biological Process | T2R | 4 | 0.034417 |
| GO:0035556~intracellular signal transduction | Biological Process | T2R | 4 | 0.034417 |
| GO:0044700~single organism signalling | Biological Process | T2R | 4 | 0.034417 |
| GO:0034470~ncRNA processing | Biological Process | T2R | 6 | 0.046209 |

\*Count indicates the number of genes in each GO term class. These are the genes that has a $\log_2FC$ of $>0.6$ .

**Table 3:** The log_2_FC for DEGs among the antigen coding genes in IEDB database for *M. leprae*

| Gene name | FC_NR | FC_T1R | FC_T2R | p-value_T1R | p-value_T2R |
| --- | --- | --- | --- | --- | --- |
| ML0008 | 0.060 | 1.220 | 0.830 | 0.0157 | 0.0667 |
| ML0050 | 0.210 | 0.420 | 1.100 | 0.6053 | 0.2289 |
| ML0091 | -0.020 | -0.270 | -0.140 | 0.2622 | 0.2371 |
| ML0098 | 0.020 | -0.430 | -0.160 | 0.1750 | 0.5904 |
| ML0126 | 0.100 | 0.310 | 0.390 | 0.0792 | 0.1099 |
| ML0136 | 0.030 | 0.470 | 0.530 | 0.3796 | 0.2864 |
| ML0176 | 0.020 | -0.230 | -0.400 | 0.2519 | 0.2946 |
| ML0234 | -0.340 | -0.280 | -0.160 | 0.5852 | 0.6195 |
| ML0308 | 0.020 | 0.010 | -0.040 | 0.2947 | 0.1061 |
| ML0317 | 0.070 | 0.130 | 0.010 | 0.5024 | 0.0667 |
| ML0375 | -0.030 | 0.300 | 0.280 | 0.4125 | 0.1388 |
| ML0380 | 0.080 | -0.050 | 0.630 | 0.4594 | 0.2821 |
| ML0394 | 0.170 | 1.190 | 1.390 | 0.1120 | 0.0198 |
| ML0398 | 0.010 | -0.110 | -0.020 | 0.2699 | 0.3885 |
| ML0411 | 0.010 | -0.160 | -0.010 | 0.5904 | 0.7327 |
| ML0576 | 0.080 | 0.150 | 0.020 | 0.1918 | 0.2723 |
| ML0611 | 0.000 | -0.080 | -0.230 | 0.1740 | 0.2657 |
| ML0638 | 0.060 | -0.350 | -0.460 | 0.1855 | 0.0651 |
| ML0726 | 0.000 | -0.030 | -0.280 | 0.0816 | 0.1436 |
| ML0757 | 0.060 | 1.330 | 0.940 | 0.0006 | 0.1158 |
| ML0840 | -0.030 | -1.030 | -1.160 | 0.0028 | 0.0004 |
| ML0841 | 0.020 | 0.570 | 0.420 | 0.0792 | 0.3072 |
| ML0885 | 0.040 | 0.360 | 0.560 | 0.0572 | 0.0869 |
| ML0987 | -0.070 | -0.590 | -0.650 | 0.1609 | 0.0398 |
| ML1057 | 0.040 | 0.140 | -0.020 | 0.1972 | 0.2291 |
| ML1189 | 0.040 | -0.280 | -0.050 | 0.4162 | 0.4799 |
| ML1207 | -0.080 | -0.460 | -0.680 | 0.0641 | 0.0671 |
| ML1214 | 0.030 | 0.370 | 0.250 | 0.2873 | 0.3489 |
| ML1217 | -0.020 | 0.260 | 0.250 | 0.2007 | 0.1549 |
| ML1274 | -0.080 | 0.840 | 0.960 | 0.0265 | 0.0683 |
| ML1358 | -0.030 | 0.540 | 0.470 | 0.1187 | 0.1509 |
| ML1419 | -0.060 | -0.480 | -0.670 | 0.4055 | 0.0604 |
| ML1420 | 0.000 | -0.150 | -0.140 | 0.2311 | 0.2457 |
| ML1553 | -0.060 | -0.720 | -0.740 | 0.0067 | 0.0146 |
| ML1601 | -0.020 | -0.150 | 0.000 | 0.2092 | 0.3392 |
| ML1795 | 0.070 | 1.060 | 0.970 | 0.0258 | 0.0594 |
| ML1811 | 0.100 | 0.000 | -0.090 | 0.3448 | 0.3400 |
| ML1812 | 0.010 | -0.130 | 0.320 | 0.4159 | 0.4812 |
| ML1829 | 0.000 | 0.310 | 0.390 | 0.3728 | 0.1712 |
| ML1891 | -0.070 | 0.070 | 0.160 | 0.3920 | 0.4598 |
| ML1915 | 0.010 | 0.070 | 0.090 | 0.1735 | 0.1664 |
| ML1923 | -0.050 | -0.880 | -0.950 | 0.0004 | 0.0532 |
| ML1989 | 0.000 | 0.410 | 0.510 | 0.0749 | 0.0581 |
| ML1990 | -1.060 | 0.690 | 1.400 | 0.2225 | 0.2180 |
| ML2028 | -0.050 | -0.400 | -0.480 | 0.0254 | 0.0399 |
| ML2038 | 0.000 | -0.430 | -0.550 | 0.0787 | 0.0299 |
| ML2055 | 0.060 | 0.320 | 0.890 | 0.2013 | 0.0559 |
| ML2069 | -0.010 | 0.090 | 0.010 | 0.3061 | 0.1957 |
| ML2283 | 0.290 | 0.470 | -0.210 | 0.4902 | 0.3048 |
| ML2347 | 0.040 | -0.300 | 0.100 | 0.2942 | 0.0083 |
| ML2395 | 0.110 | 0.440 | 0.280 | 0.3971 | 0.5247 |
| ML2400 | 0.040 | 0.240 | 0.330 | 0.1476 | 0.1593 |
| ML2452 | -0.020 | 0.520 | 0.490 | 0.0712 | 0.0858 |
| ML2496 | -0.120 | 0.040 | -0.070 | 0.3343 | 0.1724 |
| ML2531 | 0.260 | 0.610 | 0.490 | 0.6400 | 0.4597 |
| ML2535 | -0.340 | -2.490 | -3.260 | 0.0513 | 0.0092 |
| ML2567 | 0.060 | -1.880 | -1.990 | 0.0525 | 0.0245 |
| ML2688 | -0.020 | 0.020 | 0.100 | 0.3925 | 0.4782 |

**Table 4:**
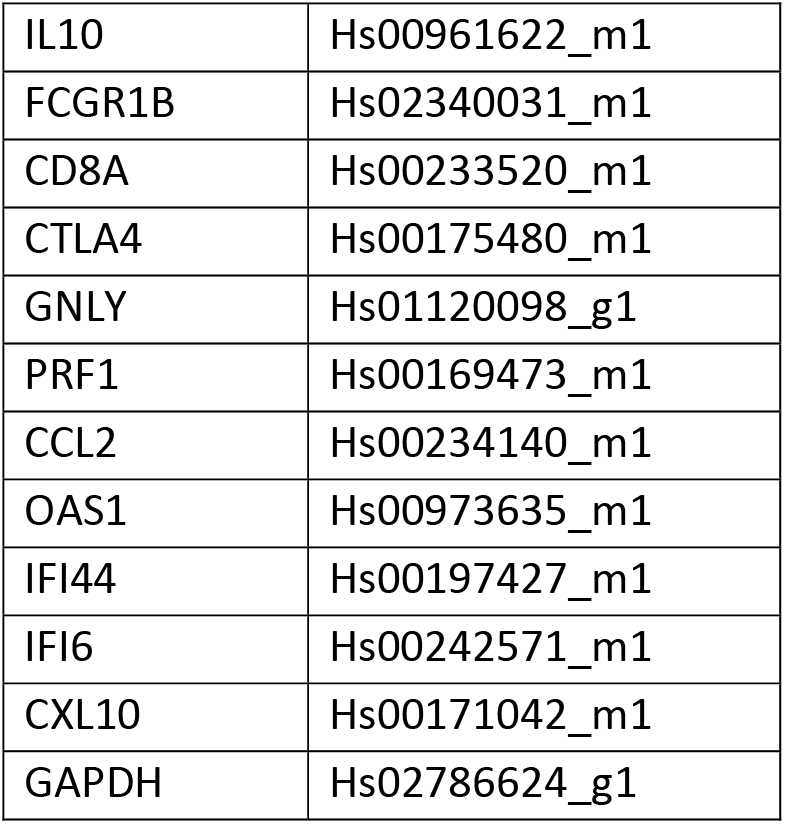
List of assays and primers

### Statistical Analysis

For the transcriptome data, normalization of DEGs and the expression threshold of + 0.6 for log_2_fold_change (log_2_FC) was determined using Agilent Gene Spring GX (Agilent Inc.). The principal component analysis was performed using in-built prcomp () function in R and plots were generated using ggplot2 package in R. The functional GO term enrichment analysis was performed using DAVID database and software.

## RESULTS

### Analysis of Transcriptome-wide changes using Principal Component Analysis

Considering 1600 genes and 45 rRNA transcripts whose intensities were noted from the hybridization arrays, the normalized data with log_2_FC values were subjected to principal component (PC) analysis to estimate the sample variance and reduce the dimensionality in the data. We first determined if the number of PCs are sufficient to explain the fraction of variance using a Pareto chart. A sequential reduction in variance across PCs was noted with PC 53 being zero indicating that the sample numbers are sufficient to explain variance. Further we visualized the clusters using ggfortify () package in R and plotted the clusters from the PCs. Fig: 2A &2B.

**Fig 2.**
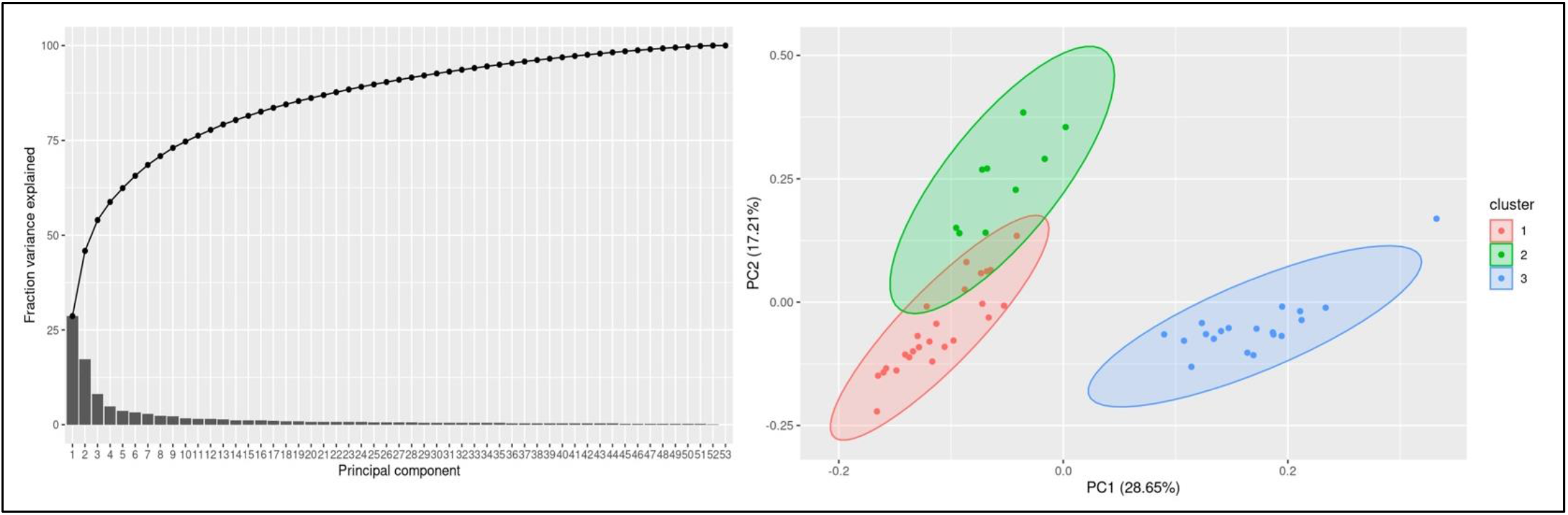
**A:** A pareto plot with fraction variance explained using PCs for each of the sample and **2B** with clusters mapped between PC1 and PC2.

Cluster 1 represents T1R, Cluster 2 represents NR, Cluster 3 represents T2R in the above figure 2A.

### DEGs of *M. leprae* across the reactional states

From the log_2_FC values, we noted transcripts corresponding to 132 genes of *M. leprae* for T1R and 117 genes in T2R that are significantly upregulated in comparison to NR. In both the reactional states, 70 genes were upregulated, and 38 genes were downregulated.. Benjamin Hochberg adjusted *p-values* were *<0.05* for all these associations. We identified only one gene (ML2664) that was upregulated in T1R and downregulated in T2R and no DEGs in converse. (Supplementary Material 1). The top 10 upregulated and the lower 10 downregulated genes were labelled in the volcano plots in fig 3B & C.

**Fig 3:**
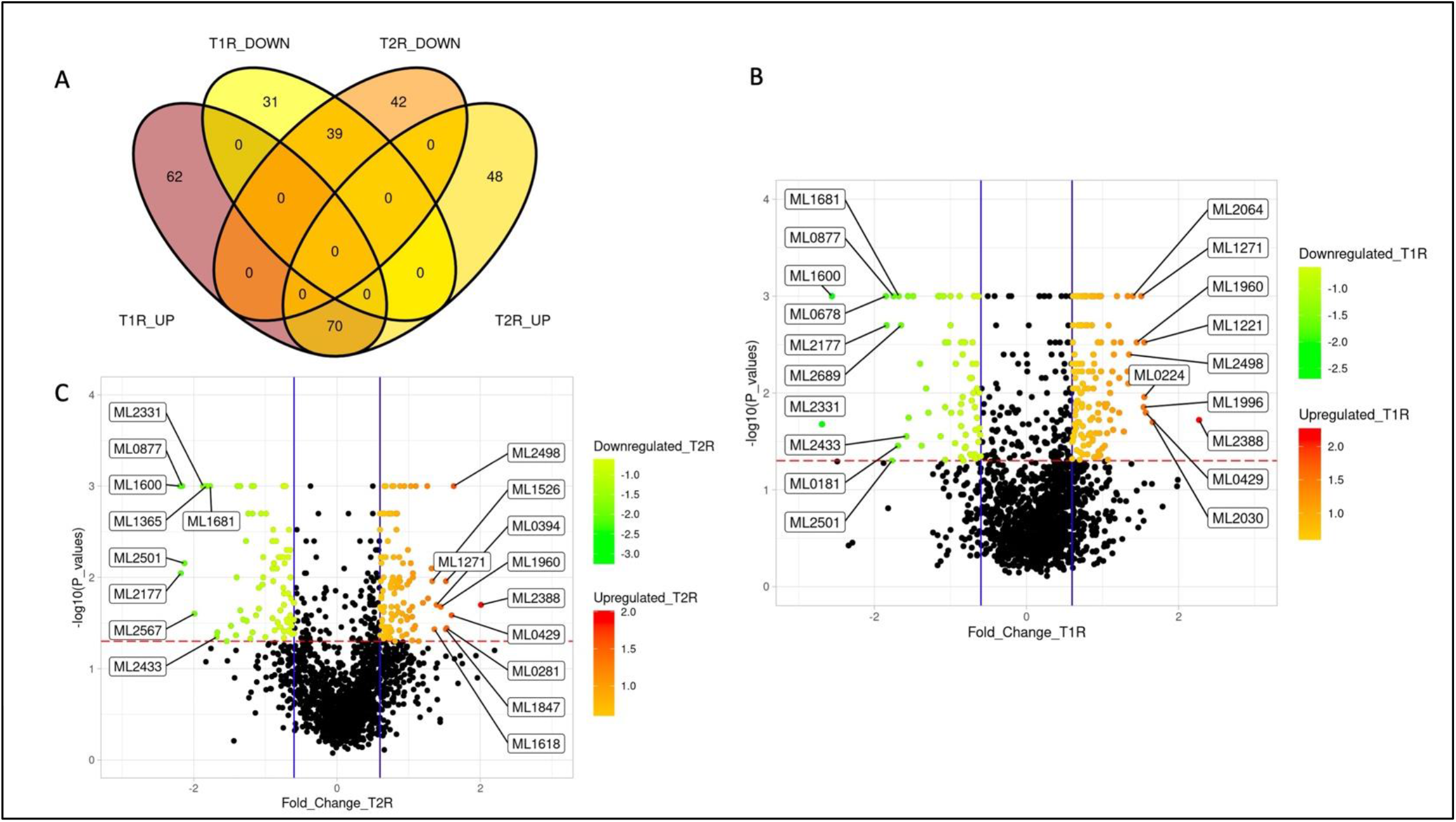
A: Venn diagram representing the number of significantly upregulated and downregulated genes of *M. leprae* across the study groups. B & C: Volcano plots with log_2_FC (derived from normalized gene expression data) showing the top 10 differentially expressed genes in T1R and T2R compared to Non reactional cases.

### Unsupervised clustering analysis for expression patterns

Hierarchical clustering analysis with the z scores of the fold changes gene-wise across the study groups revealed clusters with various enriched GO terms. Among the upregulated genes in T1R, we noted over representation of genes that encode integral membrane proteins, followed by cytosolic and ribosome bound protein coding genes (Fig 4A). From the GO biological processes genes corresponding to proteins that mediate cell wall biosynthesis, lipid biosynthetic pathways, drug transport, fatty acid & amino acid metabolism, and the biotin & folic acid biosynthetic pathways are overrepresented (Fig 4B). In the T2R, among the genes that had GO annotations for cellular component, those that encode integral membrane components were higher in number (Fig 4C) and from the GO biological processes (Fig 4D), genes involved in translation, DNA recombination, fatty acid metabolism, protein transport and aminoacid metabolism are present.

**Fig 4:**
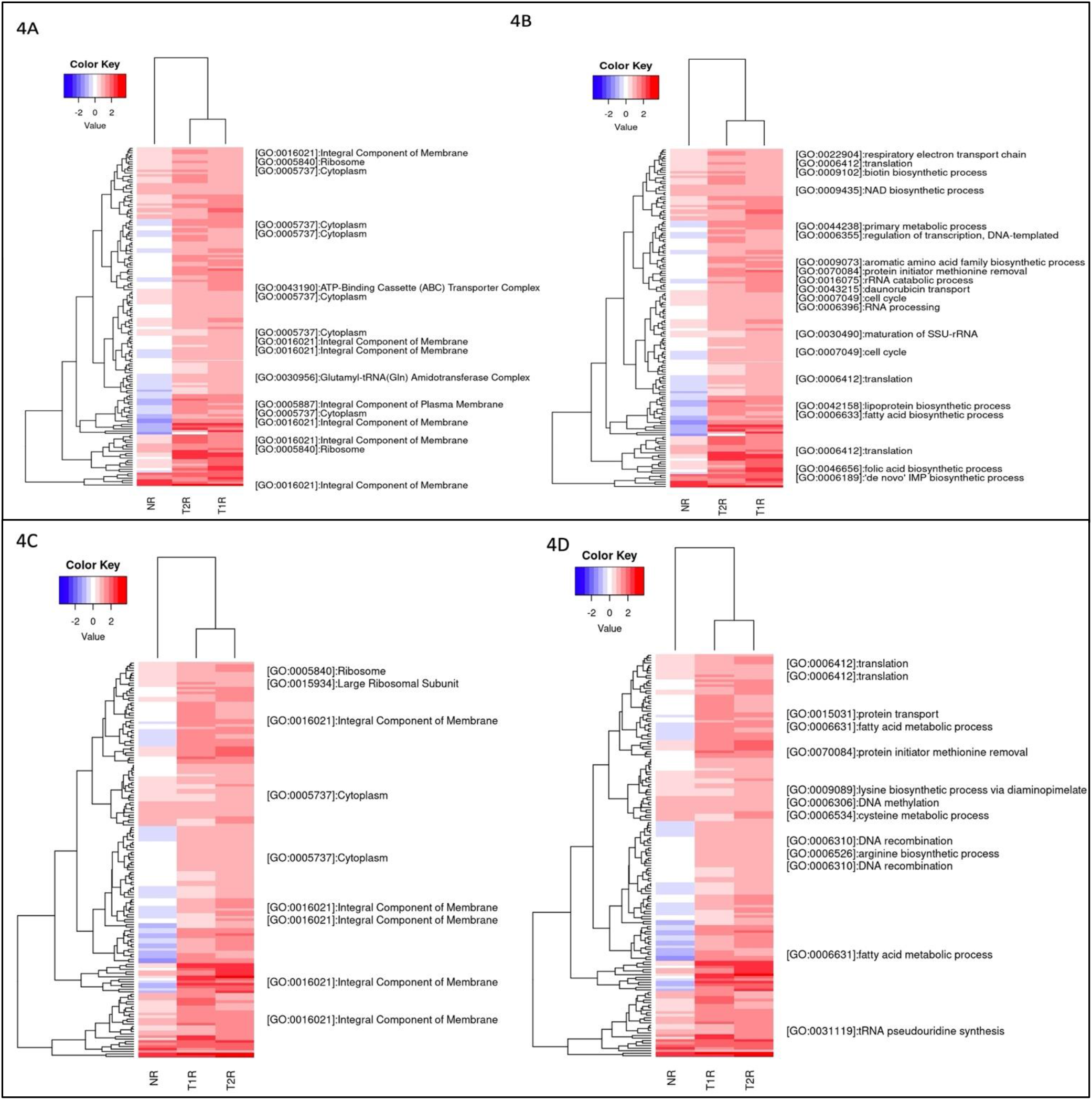
Heatmap representing significantly upregulated genes in T1R in comparison to T2R and NR and characterized by GO Cellular Component (A) and biological processes (B). Similarly, for T2R in comparison to T1R and NR, the upregulated genes with GO Cellular component (C) and biological processes were characterized (D). The color key represents normalized log2FC. The blank lines in GO terms are genes without GO annotations.

### Functional Term Analysis from GO Annotations

We further used the enriched GO terms from the DAVID database to ascertain the probability of gene co-occurrence in each GO term across the sample. Among the upregulated genes in T1R, for GO cellular component terms for integral component of membrane [GO:0016021], intrinsic component of membrane [GO:0031224], the membrane part [GO:0044425] and membrane [GO:0016020] were significantly overrepresented (*P-value 0.0004*). In the GO biological processes, terms for transcription and regulation of transcription, RNA biogenesis, regulation of RNA biosynthetic processes and lipid metabolism were overrepresented among others noted in Table2 and Fig 5A. For the GO Molecular Function, genes involved in DNA binding and hydrolase activity are noted. In T2R, for GO cellular component terms for integral component of membrane [GO:0016021], intrinsic component of membrane [GO:0031224], the membrane part [GO:0044425] and membrane [GO:0016020] were significantly overrepresented (*P-value 0.008*). For the Biological Process terms, lipid metabolic processes, cellular component biogenesis, RNA processing, RNA metabolic process, ribonucleoprotein complex biogenesis and genes involved in signal transduction were significantly overexpressed along with others shown in Table 2(Supplementary), (Fig 5B).

**Fig 5A:**
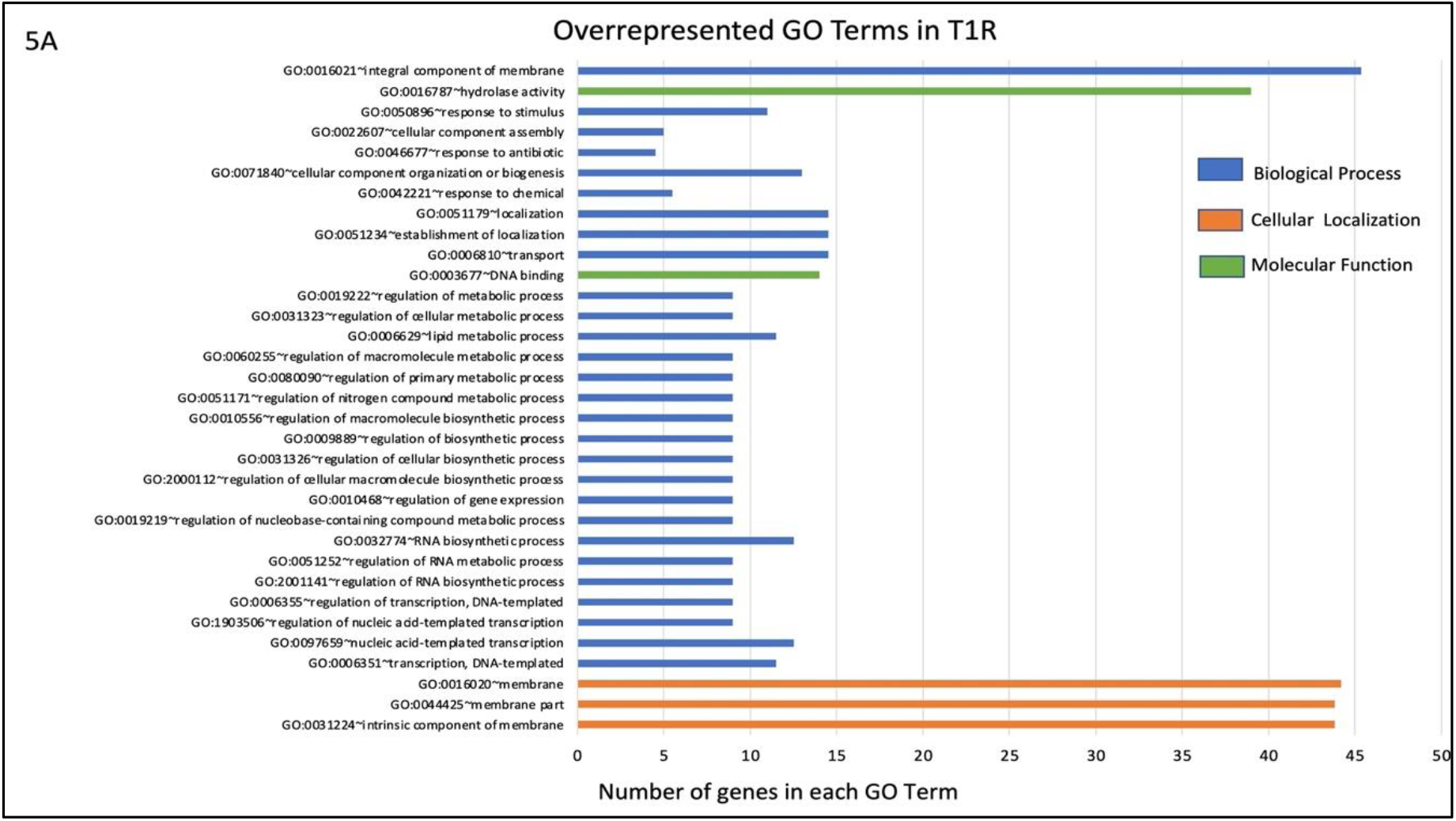
Overrepresented GO Terms in the T1R Group (*p<0.05*). The number of genes on x-axis indicates the number genes that have the same GO term in each of the classes (biological process, cellular localization and molecular function.

**Fig 5B:**
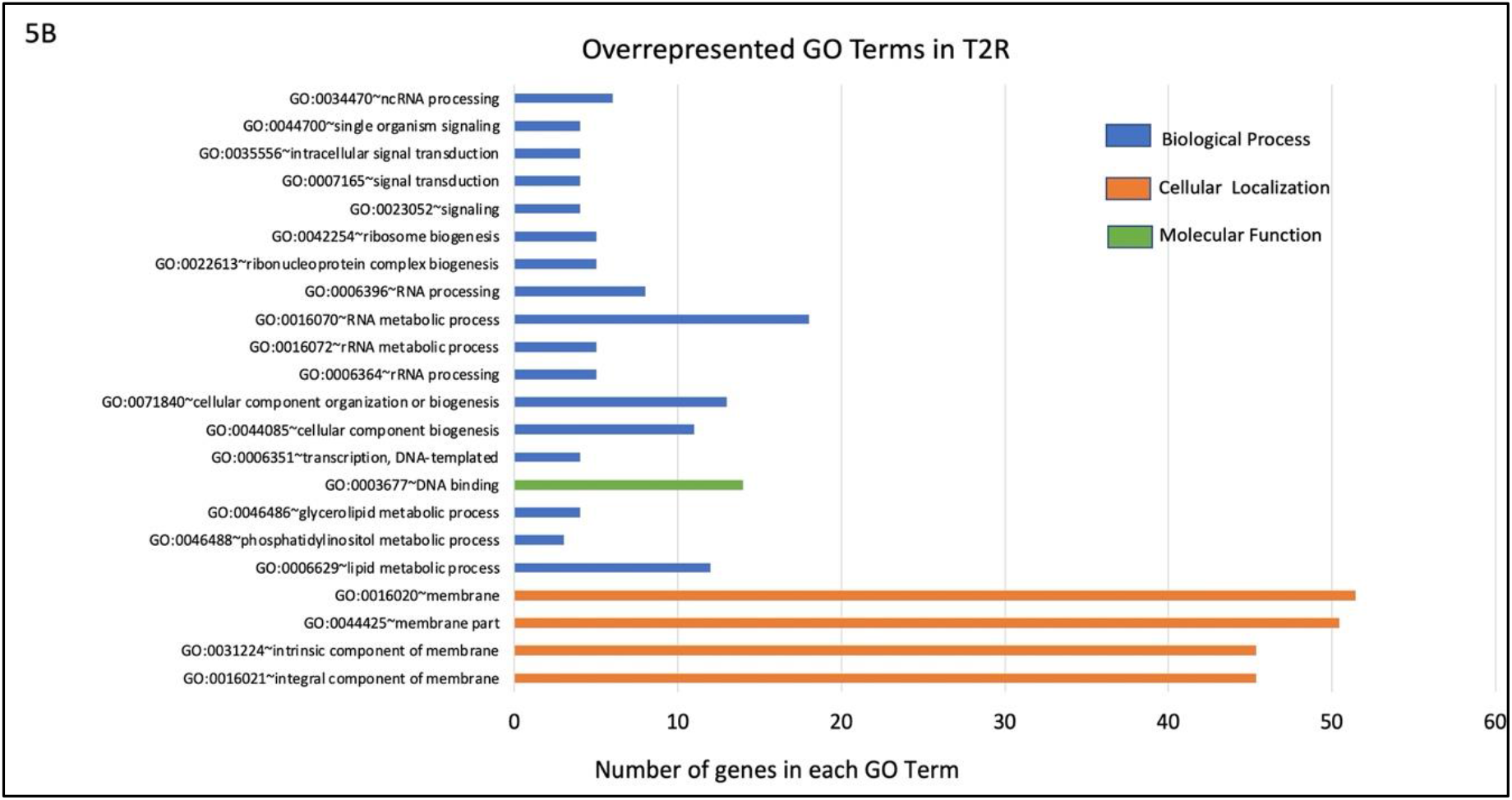
Overrepresented GO Terms in the T2R Group (*p<0.05*). The number of genes on x-axis indicates the number genes that have the same GO term in each of the classes (biological process, cellular localization and molecular function.

### Expression profiles of known *M. leprae* antigens from the IEDB database

From the 61 antigens recorded in Immune Epitope Database (IEDB) for *M. leprae* (identified by the search term *Bacillus leprae*), we selected 58 protein antigens which has Uniprot IDs (Table-3) and studied their expression across the sample. The significantly upregulated antigen coding genes in T1R and T2R are depicted in Fig 6 A and B respectively.

**Fig 6A:**
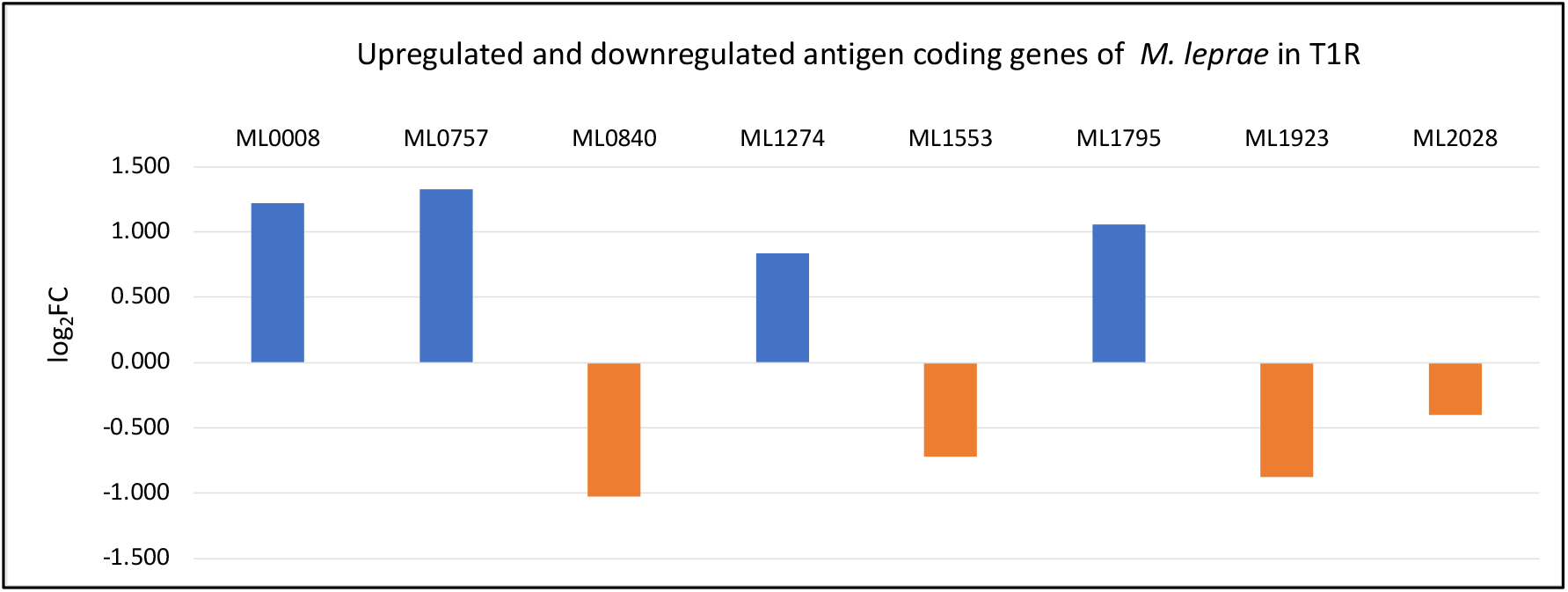
DEGs among antigen coding genes in T1R

**Fig 6B:**
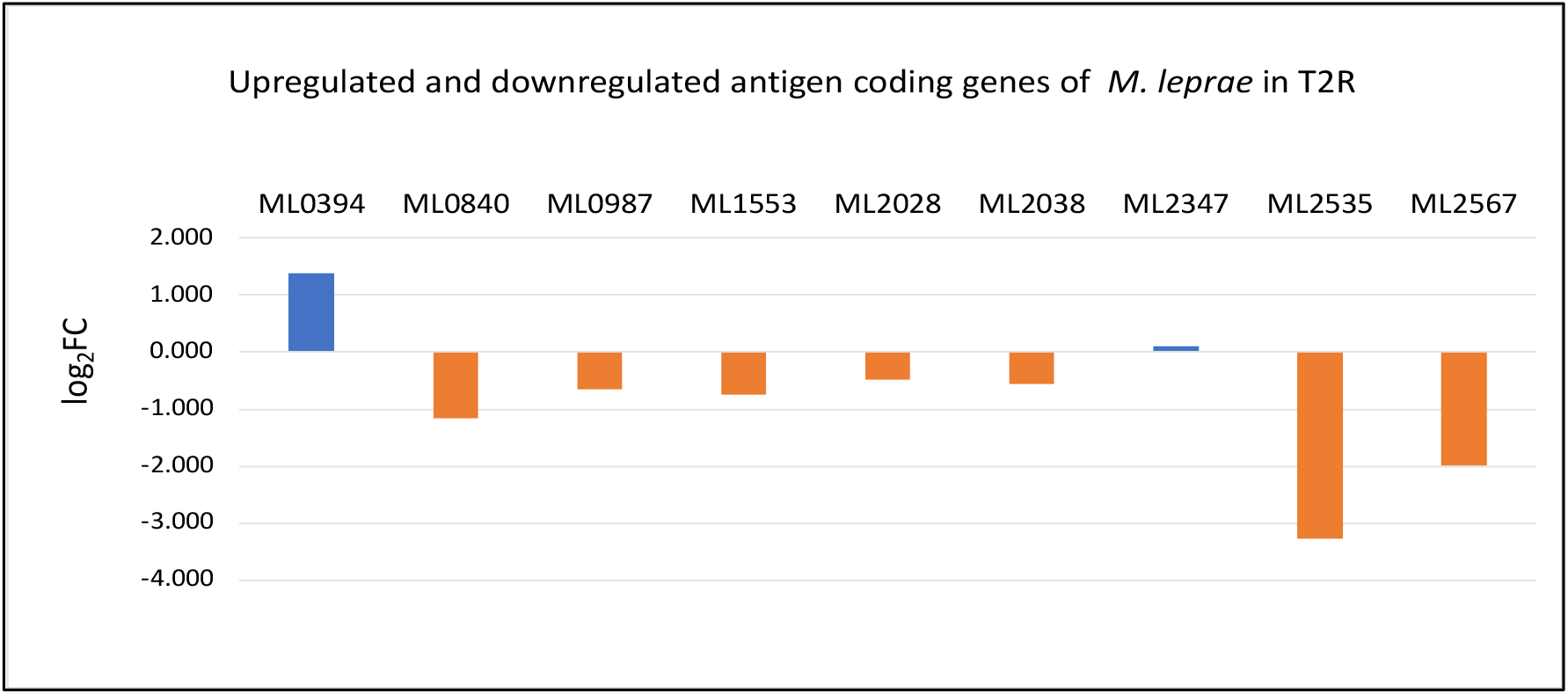
DEGs among antigen coding genes in T2R

### Consistent overexpression of ML2388 across the sample in reactional states of leprosy

Among all the significant DEGs noted across the sample, we have seen consistent over expression of ML2388 across the reactional states T1R and T2R. This gene encodes a possible membrane protein and has GO term for cellular localization as the integral component of the membrane. We predicted linear B cell epitopes on this protein using Bepipred 2.0. The results were presented in Fig 7. Three linear epitopes were predicted with high confidence in the exposed and helical or coiled regions of the protein(15).

**Fig 7:**
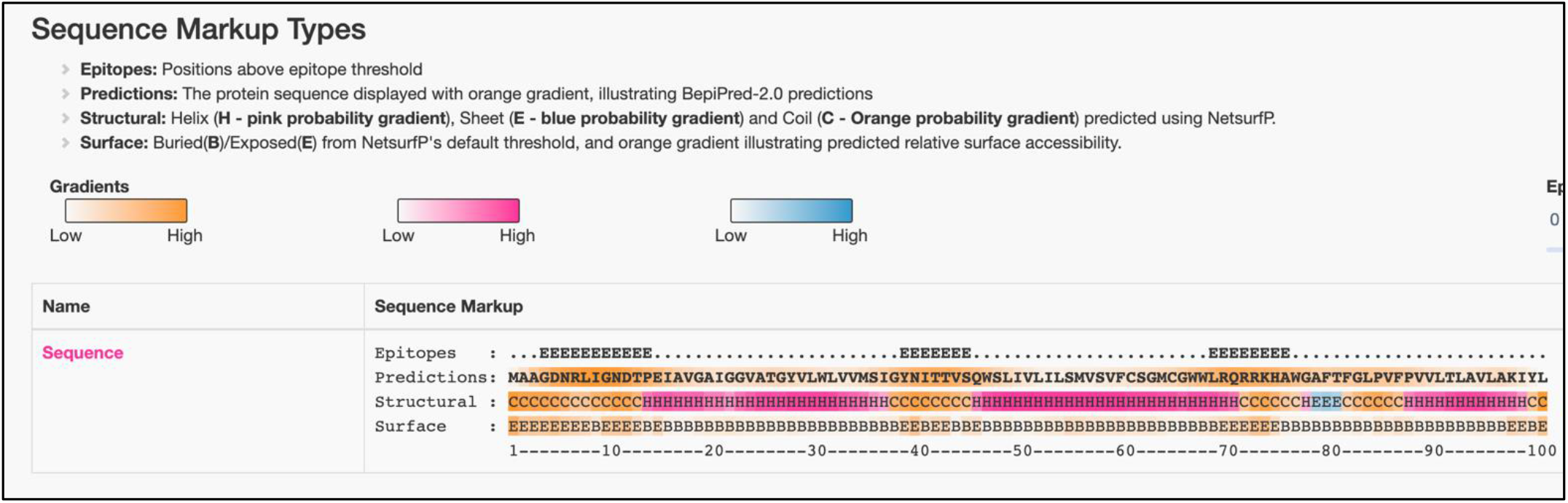
Predicted linear B-Cell Epitopes in *ML2388*

### Differential Expression of Human Immune Genes

From the RT2 PCR Profiler arrays, the expression data was analysed using Qiagen GeneGlobe pipeline using the ΔΔCt method. The threshold cycle values across the study groups were normalized using the house keeping gene (GAPDH (Glyceraldehyde-3-phosphate dehydrogenase)) expression levels (8). The quality checks were performed by measuring the PCR array reproducibility, ΔCt of the reverse transcription control and genomic DNA contamination. For the test sets (T1R and T2R), the PPC was 15 which indicates that the array reproducibility check has been passed, the transcription control has a ΔCt of 5.28 and ΔCt of genomic DNA contamination is 33 which indicates that the contamination is minimal. The cut off Ct value is set to 33. Fold-Change (2^(-ΔΔCt)^) is the normalized gene expression (2(^-ΔCt)^) in the Test Sample divided the normalized gene expression (2(^-ΔCt)^) in the Control Sample. Fold-Regulation represents fold-change results in a biologically meaningful way.

We noted a significant upregulation of CXC chemokines, CXCL9, CXCL10, CXCL2, CXCL11, CD40 ligand (CD40LG), and interleukin IL17A in T1R. In T2R, CXC chemokines CXCL10, CXCL11, CXCL9, CXCL2 and CD40 ligand (CD40LG) were upregulated (16).

### Multiplex qPCR Assays

We also tested the expression levels of a selected set of human immune genes that were observed elsewhere(17–19) to have associations with reactional states in leprosy using the multiplex qPCR system in our study groups (T1R (n=16), T2R (n=9) and NR (n=16)). Between the NR and the T1R groups, a significant difference in expression was observed for genes GNLY (Granulysin), CD8A, CXCL10, IL10, PRF1 (Perforin 1), CCL2, FCGR1B (Fc Gamma Receptor Ib), OAS1 (2’-5’-Oligoadenylate Synthetase 1), IFI44 (Interferon Induced Protein 44) and CTLA4 (cytotoxic T-lymphocyte-associated protein 4). Gene expression was higher (lower ΔCt) for all the genes tested in patients with T1R than NR patients. No statistically significant differences in expression were noted between NR and T2R or T1R and T2R.

**Fig 8:**
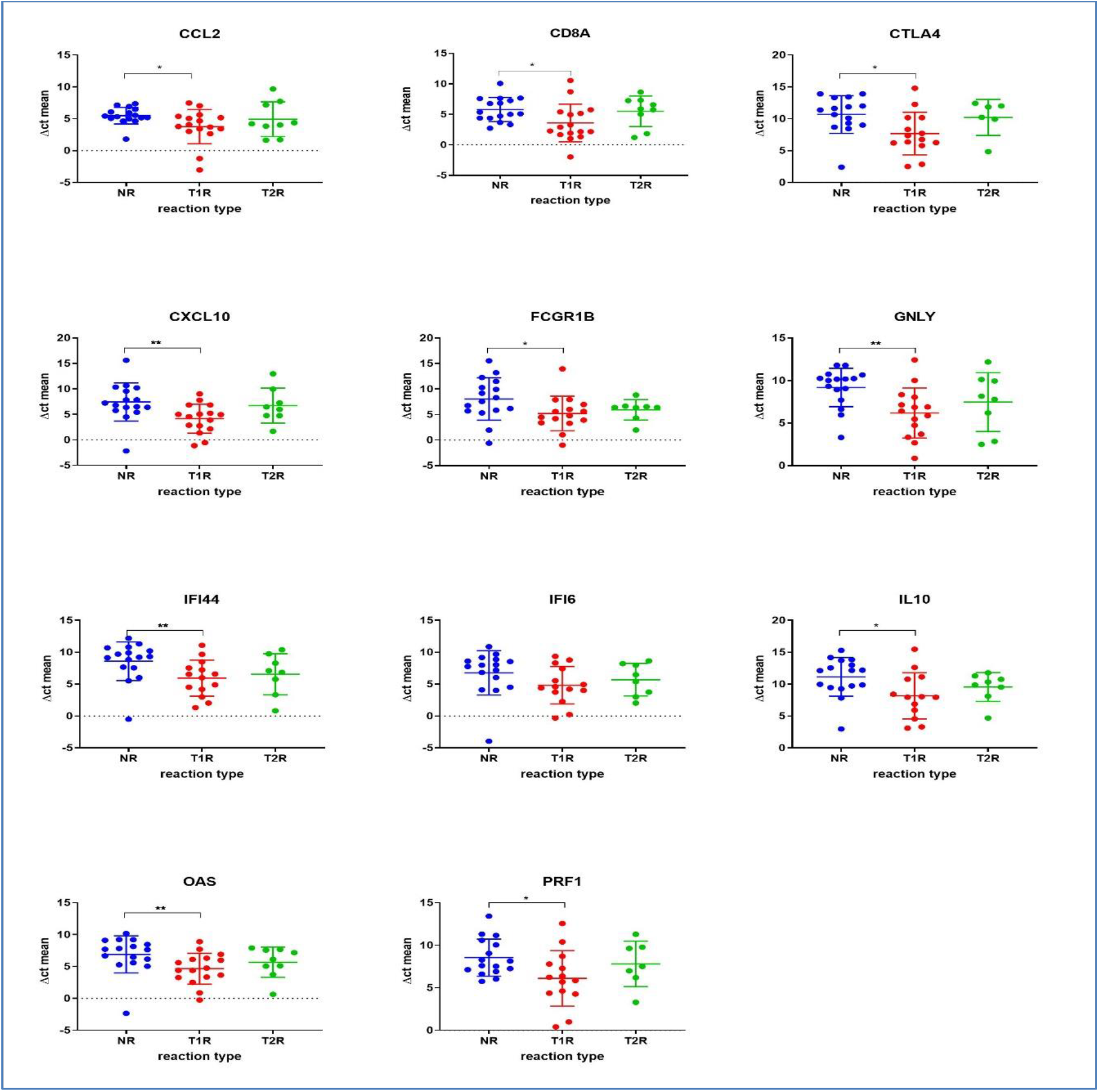
The panel depicts immune gene expressions across the study groups as determined by qPCR. Statistically significant differences in mean of the ΔCt for technical replicates of each of the sample was denoted by * for p-value of 0.05 and ** for p-value of *0.005*.

## Discussion

Reactional states in leprosy pose a significant challenge to the global efforts to contain leprosy. These immune exacerbations are a major cause of nerve function impairment and consequent disabilities in leprosy. Various host-related factors have been reported as risk factors for Type 1 reactions these include increasing age, extensive disease, and having a positive slit-skin smear. LL and a bacillary index greater than 4+ are established risk factors for Type 2 reactions(20). Few studies have also implicated a hypothesis of antigenic triggers in Type 1 Reaction, leading to expansion of both cross-reactive and specific T-cells. Studies also says that little genotypic variation exists between strains of M. leprae, a fact inconsistent with the high degree of variability in virulence and disease penetrance between individuals. This suggests that success of infection and leprosy progression rests in large part upon the host’s immune response and genetic complement(21). Various studies *M. leprae* and host immune gene expression signatures can be potential biomarkers for diagnostic test development. In this study, we conducted a case-control association analysis of DEGs in *M. leprae* transcriptome in the lesional skin tissues of leprosy cases in T1R and T2R and compared them with those without any hypersensitivity reactions to decipher a gene expression signature for these reactional states. From the GEO datasets for *M. leprae*, while there are several studies on understanding host transcriptomic response(18,22–24) to *M. leprae* infection, we identified only one other study (25) that profiled differential gene expression within *M. leprae* transcriptome to ascertain their implications on reactional states. Although, our experiments were conducted with total RNA, we limited our analysis to expression profiles in the protein coding genes. While identifying DEGs across study groups remained the main objective, we attempted to delineate the possible functional implications of DEGs on the reactional outcomes. The gene chip array from Agilent that we employed in the experiments is available at the GEO repository (GPL22363).

From the upregulated genes in T1R, we noted 27 genes that encode integral and intrinsic components of the cell wall and plasma membrane (supplementary-table-S2). We noted 12 virulence genes among the upregulated which include ML0114 - an ABC O-antigen transporter, ML1014 – RNA polymerase sigma factor sigB, ML1076-RNA(26) polymerase sigma factor SigE, ML1128 - Diaminopimelate decarboxylase, ML1220-Biotin synthase(27), ML1547-4’-phosphopantetheinyl transferase, ML1656-3-oxoacyl-[acyl-carrier-protein] synthase, ML1675-Uracil-DNA glycosylase, ML2124-Sensor-type histidine kinase, ML2307 - Transcriptional regulator and ML2350 - ATP-dependent efflux pump essential for phthiocerol dimycocerosates translocation(28,29). Overexpression of virulence genes were noted in several other studies in reactional states (5, 6). We also noted over expression of heat shock proteins (*hsp18*) in T1R (31).

From the upregulated genes in T2R, we noted that 10 virulence genes were significantly over expressed. They are ML0243 - Putative acyl-CoA synthetase, ML1014–RNA polymerase sigma factor sigB, ML1076 - RNA polymerase sigma factor SigE, ML1128 - Diaminopimelate decarboxylase, ML1633 - Possible secreted hydrolase, ML1727 - O-phosphoserine phosphohydrolase(32), ML1925 - Superoxide dismutase [Cu-Zn], ML1954-Pantothenate kinase(33), ML2307 - Transcriptional regulator and ML2439 - Sensory transduction protein RegX3(34–36).

From the human host genes, we noted that for the NR and the T1R groups, a significant difference in expression was observed for genes GNLY (Granulysin) responsible for transfer of granzymes which are a family of serine proteases traditionally known for their role in promoting cytotoxicity of foreign, infected or neoplastic cells, CD8A (Cytotoxic T-lymphocytes) important for immune defense against intracellular pathogens, including viruses and bacteria, CXCL10 (Interferon gamma induced protein-10). Granulysin has been noted to express high in tuberculoid lesions.(37). Upregulation of CD8 antigen in reversal reactions has also been noted leprosy/HIV co-infections (38).

Many studies reveal the upregulation of CXCL10 (IP10) in reactional states of leprosy (39)(40)(41).

Expression of IP-10 is seen in many Th1-type inflammatory diseases, where it is thought to play an important role in recruiting activated T cells into sites of tissue inflammation, Interleukin 10 (IL-10), a cytokine with potent anti-inflammatory properties that plays a central role in limiting host immune response to pathogens, thereby preventing damage to the host and maintaining normal tissue homeostasis, PRF1 (Perforin 1) a glycoprotein responsible for pore formation in cell membranes of target cells., CCL2 chemokine which controls immunity by promoting regulatory T cell communication with dendritic cells in lymph nodes. FCGR1B (Fc Gamma Receptor Ib) functions ad humoral immune response by binding to the Fc regions of immunglobulin, OAS1 (2’-5’-Oligoadenylate Synthetase 1) helps in degradation of viral infections, IFI44 (Interferon Induced Protein 44) and CTLA4 (cytotoxic T-lymphocyte-associated protein 4) which acts as a halting mechanism, decreasing the function of T cells. Similar results were noted with RT2 profiler arrays in this study. Upregulation of CXC chemokines, CXCL9, CXCL10, CXCL2, CXCL11, CD40 ligand (CD40LG), and interleukin IL17A were upregulated in T1R. In T2R, CXC chemokines CXCL10, CXCL11, CXCL9, CXCL2 and CD40 ligand (CD40LG) were upregulated. (11,17,42).

In conclusion, our study profiled various upregulating and downregulating gene signatures from *Mycobacterium leprae* and human immune genes that demonstrated plausible association with reactional states in leprosy. Further studies are however required to validate the above identified gene expression signatures as predictive markers for leprosy reactions in a longitudinal cohort

## Acknowledgments

The authors would like to thank all the administration and laboratory staff at Schieffelin Institute of Health-Research and Leprosy Center who were involved in sample collection and performing the experiments. Special thanks to the administration of SIH-R&LC Karigiri for the infrastructural support throughout the study and for the financial support from the Leprosy Research Initiative (LRI) and the Turing Foundation under LRI Grant number 704.16.57.

## References

1. Richardus JH, Finlay KM, Croft RP, Smith WC. Nerve function impairment in leprosy at diagnosis and at completion of MDT: a retrospective cohort study of 786 patients in Bangladesh. Lepr Rev (1996) 67:297–305.

2. WHO Expert Committee on leprosy, World Health Organization. WHO Expert Committee on leprosy: eighth report. Geneva: World Health Organization (2012). https://apps.who.int/iris/handle/10665/75151 [Accessed March 22, 2022]

3. Ridley DS, Jopling WH. Classification of leprosy according to immunity. A five-group system. Int J Lepr Other Mycobact Dis (1966) 34:255–273.

4. Montoya D, Modlin RL. Learning from leprosy: insight into the human innate immune response. Adv Immunol (2010) 105:1–24. doi: 10.1016/S0065-2776(10)05001-7

5. Kahawita IP, Lockwood DNJ. Towards understanding the pathology of erythema nodosum leprosum. Trans R Soc Trop Med Hyg (2008) 102:329–337. doi: 10.1016/j.trstmh.2008.01.004

6. Nery JA da C, Filho FB, Quintanilha J, Machado AM, Oliveira S de SC, Sales AM. Understanding the type 1 reactional state for early diagnosis and treatment: a way to avoid disability in leprosy. An Bras Dermatol (2013) 88:787–792. doi: 10.1590/abd1806-4841.20132004

7. Sehgal VN. Reactions in leprosy. Clinical aspects. Int J Dermatol (1987) 26:278–285. doi: 10.1111/j.1365-4362.1987.tb00188.x

8. Kwittken J. Erythema nodosum leprosum. Int J Lepr Other Mycobact Dis (1968) 36:94–95.

9. Yuan Y-H, Liu J, You Y-G, Chen X-H, Yuan L-C, Wen Y, Li HY, Zhang Y. Transcriptomic Analysis of Mycobacterium leprae-Stimulated Response in Peripheral Blood Mononuclear Cells Reveal Potential Biomarkers for Early Diagnosis of Leprosy. Front Cell Infect Microbiol (2021) 11:714396. doi: 10.3389/fcimb.2021.714396

10. Leal-Calvo T, Avanzi C, Mendes MA, Benjak A, Busso P, Pinheiro RO, Sarno EN, Cole ST, Moraes MO. A new paradigm for leprosy diagnosis based on host gene expression. PLoS Pathog (2021) 17:e1009972. doi: 10.1371/journal.ppat.1009972

11. Tió-Coma M, Kiełbasa SM, van den Eeden SJF, Mei H, Roy JC, Wallinga J, Khatun M, Soren S, Chowdhury AS, Alam K, et al. Blood RNA signature RISK4LEP predicts leprosy years before clinical onset. EBioMedicine (2021) 68:103379. doi: 10.1016/j.ebiom.2021.103379

12. Rajkumar AP, Qvist P, Lazarus R, Lescai F, Ju J, Nyegaard M, Mors O, Børglum AD, Li Q, Christensen JH. Experimental validation of methods for differential gene expression analysis and sample pooling in RNA-seq. BMC Genomics (2015) 16:548. doi: 10.1186/s12864-015-1767-y

13. Tió-Coma M, Kiełbasa SM, van den Eeden SJF, Mei H, Roy JC, Wallinga J, Khatun M, Soren S, Chowdhury AS, Alam K, et al. Blood RNA signature RISK4LEP predicts leprosy years before clinical onset. EBioMedicine (2021) 68:103379. doi: 10.1016/j.ebiom.2021.103379

14. Goeman JJ, van de Geer SA, de Kort F, van Houwelingen HC. A global test for groups of genes: testing association with a clinical outcome. Bioinformatics (2004) 20:93–99. doi: 10.1093/bioinformatics/btg382

15. Reece ST, Ireton G, Mohamath R, Guderian J, Goto W, Gelber R, Groathouse N, Spencer J, Brennan P, Reed SG. ML0405 and ML2331 are antigens of Mycobacterium leprae with potential for diagnosis of leprosy. Clin Vaccine Immunol (2006) 13:333–340. doi: 10.1128/CVI.13.3.333-340.2006

16. Scollard DM, Chaduvula MV, Martinez A, Fowlkes N, Nath I, Stryjewska BM, Kearney MT, Williams DL. Increased CXC Ligand 10 Levels and Gene Expression in Type 1 Leprosy Reactions ▿. Clin Vaccine Immunol (2011) 18:947–953. doi: 10.1128/CVI.00042-11

17. Geluk A, van Meijgaarden KE, Wilson L, Bobosha K, van der Ploeg-van Schip JJ, van den Eeden SJF, Quinten E, Dijkman K, Franken KLMC, Haisma EM, et al. Longitudinal immune responses and gene expression profiles in type 1 leprosy reactions. J Clin Immunol (2014) 34:245–255. doi: 10.1007/s10875-013-9979-x

18. Teles RMB, Graeber TG, Krutzik SR, Montoya D, Schenk M, Lee DJ, Komisopoulou E, Kelly-Scumpia K, Chun R, Iyer SS, et al. Type I interferon suppresses type II interferon-triggered human anti-mycobacterial responses. Science (2013) 339:1448–1453. doi: 10.1126/science.1233665

19. Tió-Coma M, van Hooij A, Bobosha K, van der Ploeg-van Schip JJ, Banu S, Khadge S, Thapa P, Kunwar CB, Goulart IM, Bekele Y, et al. Whole blood RNA signatures in leprosy patients identify reversal reactions before clinical onset: a prospective, multicenter study. Sci Rep (2019) 9:17931. doi: 10.1038/s41598-019-54213-y

20. New insights in the pathogenesis of type 1 and type 2 lepra reaction. Indian Journal of Dermatology, Venereology and Leprology (2013) https://ijdvl.com/new-insights-in-the-pathogenesis-of-type-1-and-type-2-lepra-reaction/ [Accessed June 16, 2022]

21. Texereau J, Chiche J-D, Taylor W, Choukroun G, Comba B, Mira J-P. The importance of Toll-like receptor 2 polymorphisms in severe infections. Clin Infect Dis (2005) 41 Suppl 7:S408–415. doi: 10.1086/431990

22. Zhang X, Cheng Y, Zhang Q, Wang X, Lin Y, Yang C, Sun J, Huang H, Li Y, Sheng Y, et al. Meta-Analysis Identifies Major Histocompatiblity Complex Loci in or Near HLA-DRB1, HLA-DQA1, HLA-C as Associated with Leprosy in Chinese Han Population. J Invest Dermatol (2019) 139:957–960. doi: 10.1016/j.jid.2018.09.029

23. Leal-Calvo T, Martins BL, Bertoluci DF, Rosa PS, de Camargo RM, Germano GV, Brito de Souza VN, Pereira Latini AC, Moraes MO. Large-Scale Gene Expression Signatures Reveal a Microbicidal Pattern of Activation in Mycobacterium leprae-Infected Monocyte-Derived Macrophages With Low Multiplicity of Infection. Front Immunol (2021) 12:647832. doi: 10.3389/fimmu.2021.647832

24. Manry J, Nédélec Y, Fava VM, Cobat A, Orlova M, Thuc NV, Thai VH, Laval G, Barreiro LB, Schurr E. Deciphering the genetic control of gene expression following Mycobacterium leprae antigen stimulation. PLoS Genet (2017) 13:e1006952. doi: 10.1371/journal.pgen.1006952

25. Montoya DJ, Andrade P, Silva BJA, Teles RMB, Ma F, Bryson B, Sadanand S, Noel T, Lu J, Sarno E, et al. Dual RNA-Seq of Human Leprosy Lesions Identifies Bacterial Determinants Linked to Host Immune Response. Cell Rep (2019) 26:3574–3585.e3. doi: 10.1016/j.celrep.2019.02.109

26. Williams DL, Pittman TL, Deshotel M, Oby-Robinson S, Smith I, Husson R. Molecular Basis of the Defective Heat Stress Response in Mycobacterium leprae. J Bacteriol (2007) 189:8818–8827. doi: 10.1128/JB.00601-07

27. Lastória JC, de Abreu MAMM. Leprosy: a review of laboratory and therapeutic aspects - Part 2. An Bras Dermatol (2014) 89:389–401. doi: 10.1590/abd1806-4841.20142460

28. Sharma R, Lavania M, Chauhan DS, Katoch K, Amresh null, Pramod null, Rakhi null, Richa null, Katoch VM. Potential of a metabolic gene (accA3) of M. leprae as a marker for leprosy reactions. Indian J Lepr (2009) 81:141–148.

29. Orlova M, Cobat A, Huong NT, Ba NN, Van Thuc N, Spencer J, Nédélec Y, Barreiro L, Thai VH, Abel L, et al. Gene set signature of reversal reaction type I in leprosy patients. PLoS Genet (2013) 9:e1003624. doi: 10.1371/journal.pgen.1003624

30. Williams DL, Torrero M, Wheeler PR, Truman RW, Yoder M, Morrison N, Bishai WR, Gillis TP. Biological implications of Mycobacterium leprae gene expression during infection. J Mol Microbiol Biotechnol (2004) 8:58–72. doi: 10.1159/000082081

31. Lini N, Shankernarayan NP, Dharmalingam K. Quantitative real-time PCR analysis of Mycobacterium leprae DNA and mRNA in human biopsy material from leprosy and reactional cases. J Med Microbiol (2009) 58:753–759. doi: 10.1099/jmm.0.007252-0

32. Acebrón-García-de-Eulate M, Blundell TL, Vedithi SC. Strategies for drug target identification in Mycobacterium leprae. Drug Discovery Today (2021) 26:1569–1573. doi: 10.1016/j.drudis.2021.03.026

33. Hasan Z, Mahmood A, Zafar S, Khan AA, Hussain R. Leprosy patients with lepromatous disease have an up-regulated IL-8 response that is unlinked to TNF-alpha responses. Int J Lepr Other Mycobact Dis (2004) 72:35–44. doi: 10.1489/1544-581X(2004)072<0035:LPWLDH>2.0.CO;2

34. Duthie MS, Goto W, Ireton GC, Reece ST, Cardoso LPV, Martelli CMT, Stefani MMA, Nakatani M, de Jesus RC, Netto EM, et al. Use of Protein Antigens for Early Serological Diagnosis of Leprosy. Clin Vaccine Immunol (2007) 14:1400–1408. doi: 10.1128/CVI.00299-07

35. Tang ASO, Wong QY, Yeo ST, Ting IPL, Lee JTH, Fam TL, Chew LP, Chua HH, Muniandy P. Challenges in Managing a Lepromatous Leprosy Patient Complicated with Melioidosis Infection, Dapsone-Induced Methemoglobinemia, Hemolytic Anemia, and Lepra Reaction. Am J Case Rep (2021) 22:e931655. doi: 10.12659/AJCR.931655

36. Sapkota BR, Macdonald M, Berrington WR, Misch EA, Ranjit C, Siddiqui MR, Kaplan G, Hawn TR. Association of TNF, MBL, and VDR Polymorphisms with Leprosy Phenotypes. Hum Immunol (2010) 71:992–998. doi: 10.1016/j.humimm.2010.07.001

37. Geluk A, van Meijgaarden KE, Wilson L, Bobosha K, van der Ploeg-van Schip JJ, van den Eeden SJF, Quinten E, Dijkman K, Franken KLMC, Haisma EM, et al. Longitudinal Immune Responses and Gene Expression Profiles in Type 1 Leprosy Reactions. J Clin Immunol (2014) 34:245–255. doi: 10.1007/s10875-013-9979-x

38. de Oliveira AL, Amadeu TP, de França Gomes AC, Menezes VM, da Costa Nery JA, Pinheiro RO, Sarno EN. Role of CD8+ T cells in triggering reversal reaction in HIV/leprosy patients. Immunology (2013) 140:47–60. doi: 10.1111/imm.12108

39. Ferreira H, Mendes MA, de Mattos Barbosa MG, de Oliveira EB, Sales AM, Moraes MO, Sarno EN, Pinheiro RO. Potential Role of CXCL10 in Monitoring Response to Treatment in Leprosy Patients. Front Immunol (2021) 12:662307. doi: 10.3389/fimmu.2021.662307

40. van Hooij A, Tjon Kon Fat EM, Richardus R, van den Eeden SJF, Wilson L, de Dood CJ, Faber R, Alam K, Richardus JH, Corstjens PLAM, et al. Quantitative lateral flow strip assays as User-Friendly Tools To Detect Biomarker Profiles For Leprosy. Sci Rep (2016) 6:34260. doi: 10.1038/srep34260

41. Chaitanya VS, Lavania M, Nigam A, Turankar RP, Singh I, Horo I, Sengupta U, Jadhav RS. Cortisol and proinflammatory cytokine profiles in type 1 (reversal) reactions of leprosy. Immunology Letters (2013) 156:159–167. doi: 10.1016/j.imlet.2013.10.008

42. Teles RMB, Lu J, Tió-Coma M, Goulart IMB, Banu S, Hagge D, Bobosha K, Ottenhoff THM, Pellegrini M, Geluk A, et al. Identification of a systemic interferon-γ inducible antimicrobial gene signature in leprosy patients undergoing reversal reaction. PLoS Negl Trop Dis (2019) 13:e0007764. doi: 10.1371/journal.pntd.0007764

43. Geluk A, Bobosha K, van der Ploeg-van Schip JJ, Spencer JS, Banu S, Martins MVSB, Cho S-N, Franken KLMC, Kim HJ, Bekele Y, et al. New biomarkers with relevance to leprosy diagnosis applicable in areas hyperendemic for leprosy. J Immunol (2012) 188:4782–4791. doi: 10.4049/jimmunol.1103452

